# Isolator: accurate and stable analysis of isoform-level expression in RNA-Seq experiments

**DOI:** 10.1101/088765

**Authors:** Daniel C. Jones, Kavitha T. Kuppusamy, Nathan J. Palpant, Xinxia Peng, Charles E. Murry, Hannele Ruohola-Baker, Walter L. Ruzzo

## Abstract

While RNA-Seq has enabled great progress towards the goal of wide-scale isoform-level mRNA quantification, short reads have limitations when resolving complex or similar sets of isoforms. As a result, estimates of isoform abundance carry far more uncertainty than those made at the gene level. When confronted with this uncertainty, commonly used methods produce estimates that are often high-variance—small perturbations in the data often produce dramatically different results, confounding downstream analysis. We introduce a new method, Isolator, which analyzes all samples in an experiment in unison using a simple Bayesian hierarchical model. Combined with aggressive bias correction, it produces estimates that are simultaneously accurate and show high agreement between samples. In a comprehensive comparison of accuracy and variance, we show that this property is unique to Isolator. We further demonstrate that the approach of modeling an entire experiment enables new analyses, which we demonstrate by examining splicing monotonicity across several time points in the development of human cardiomyocyte cells.

## 1 Introduction

Since its introduction, RNA-Seq has rapidly become a preferred method of studying gene expression, having proved to be reliable, reproducible, and increasingly affordable. In principle, RNA-Seq enables measurements of expression at a scale and resolution than has previously been impractical or impossible. Rather than relying on an often vaguely defined notion of a gene, expression of specific isoforms or exons can be measured. In practice this promise is difficult to realize, with methods designed to detect changes in splicing rarely concuring [14].

Regardless, splicing has been repeatedly shown to be a essential regulatory mechanism, and particularly important in development and differentiation [8]. We examine alternative splicing in heart development in two contexts in Sections 2.6 and 2.7.

Most RNA-Seq procotocols produce short reads (typically 30 to 150 nucleotides, often paired-end). Because a short read is often compatible with many isoforms, the signal produced from sequencing must be deconvoluted, requiring a certain level of algorithmic sophistication. This problem was tackled early on and with great success by Cufflinks [25], and indeed many of the methods developed in the interim have followed from the same basic model of RNA-Seq in which transcripts are represented as particular distributions over possible reads, and the problem is then to infer mixing coefficients (relative expression values) inducing a mixture model that best explains the data [18].

We built on this work, implementing a new method called Isolator. Though it retains the same underlying model of RNA-Seq, we approach the problem from the perspective that isoform-level estimates possess a degree of uncertainty that renders unregularized maximum likelihood point-estimates alone a fundamentally inadequate solution to the problem. As the purpose of gene expression studies is to compare expression between multiple samples, our goal, beyond maximizing a narrow definition of accuracy, is to provide a tool that effectively accounts for this uncertainty in a coherent and reliable manner.

Rather than estimating expression of individual samples, Isolator uses a hierarchical model of an entire experiment, for example, including all samples and time points in a time series. In addition to inferring expression values for the individual samples, we introduce parameters representing condition-wise and experiment-wise expression and splicing, as well as per-transcript variance parameters that are shared across conditions. As expression studies often use only a small number of replicates, shared variance parameters allows us to make more accurate estimates of biological variability.

An efficient Markov chain Monte Carlo algorithm is used to generate samples over the parameters of this model, which are saved in an HDF5-based format [24]. The output from the sampler can then be processed to produce point estimates, credible intervals, posterior probabilities, and diagnostics. Though designed to be run on multiple samples concurrently, Isolator can be run as a conventional tool on individual samples.

The broad effect of this model is to encode the common assumption that genes tend to be similarly expressed and similarly spliced between replicates, and to a lesser extent between conditions. In the absence of sufficient evidence to the contrary, this informative prior shrinks estimates towards a baseline of no change, producing more conservative estimates of effect size than would be obtained by using, for example, a flat Dirichlet prior. This feature of the model is particularly important for robust treatment of low-expression transcripts. With few reads there is little power to determine splicing patterns. Considering the samples in isolation will often produce very different estimates. Statistical tests based on these estimates, without considering the full posterior distribution, make approximations by either disregarding the relative or compositional nature of the data and assuming independence, or by relying on asymptotic properties that may not hold when a gene is not deeply sequenced. In addition to this hierarchical model, we attempt to aggressively correct for technical effects. The naive model of RNA-Seq assumes fragments are sampled uniformly at random from transcripts in proportion to their abundance. Early on, this was shown to be far from an accurate description of real sequencing data [6]. Though severe at nucleotide resolution, gene level expression estimates are only moderately affected by this bias, as it tends to average out over long transcripts. When one considers expression at the isoform level, attempting to correct for this nonuniformity becomes vital, as the difference between two isoforms is often only a few nucleotides.

We previously developed an efficient and versatile model to account for sequence specific bias that commonly occurs at the ends of fragments [7]. In Isolator, we supplemented this with methods accounting for 3’ bias, fragment GC-content bias, and positional bias caused by end-effects of fragmentation.

Isolator is available under a permissive open source license at: https://github.com/dcjones/isolator

## 2 Results

The convention for evaluating the accuracy of RNA-Seq analysis methods has consisted chiefly of comparing point estimate accuracy in simulations, supplemented with comparisons to qPCR or other technologies. Tuning methods exclusively to maximize accuracy on simulated data risks solving only an idealized mathematical model of RNA-Seq while disregarding noise, bias, and nonuniformity. As no gold standard for whole-transcriptome isoform-level estimation exists, we adopt a multifaceted approach, focusing on measuring the variance of estimates, along with overall accuracy.

Comparisons of gene or transcript expression are sensitive to how one chooses to measure similarity. Because gene expression typically varies across many orders of magnitude, measurements like the Pearson correlation, *L*_1_ or *L*_2_ distance, or root-mean-square error are often dominated by the most highly expressed genes. Recognizing this, a popular alternative is Spearman’s rank correlation, which conversely can overemphasize the majority of transcripts, which have no, or very low expression. These issues are discussed in depth by Lovell et al [15], who propose measurements of proportionality as a more meaningful metric. We adopt their “proportionality correlation”, defined as pcor(*x, y*) = 2cov(log(*x*), log(*y*))/(var(log(*x*)) + var(log(*y*))), where is cov(*x*) and var(*x*) give the covariance and variance of *x*, respectively. Similar to other measures of correlation, proportionality correlation varies from -1 to 1, with 1 indicating perfect proportionality, and -1 perfect reciprocality. Zeros are accounted for by converting all measurements to transcripts per million (TPM) and adding 0.1. (Results were found to be fairly insensitive to the precise additive constant used here (Supplementary Section 4.1)). Though we believe proportionality correlation is a superior approach, much of this analysis was also conducted using Spearman’s rank correlation in Supplementary Section 4.2, and largely agrees with the results shown here.

Many methods have been proposed to aid in the analysis of RNA-Seq experiments. We limit our discussion specifically to those that seek to estimate the relative abundance a set of known transcripts. These include, non-exhaustively, Cufflinks [25], RSEM [12], eXpress [22], BitSeq [4], Sailfish [21], Salmon (a refinement of Sailfish currently in development), and Kallisto [1].

RSEM was used to produce both maximum likelihood and posterior mean estimates, which we label “RSEM/ML” and “RSEM/PM”, respectively. This is not an exhaustive account of such methods, but does represent a wide variety of popular approaches. The same read alignments were used for all methods other Sailfish, Salmon, and Kallisto, which are “alignment-free”, using built-in approximations to full alignment (Supplementary Section 2).

### 2.1 Agreement with qPCR and spike-in controls

Spike-in controls and qPCR are far more limited than RNA-Seq, measuring only the abundance of specific loci in the case of qPCR, or known proportions of simple artificial transcripts with spike-in controls. However, both are considered reliable enough to serve as proxy gold standards for gene-level expression estimates.

We use data generated by the Sequencing Quality Control Consortium (SEQC) [3], consisting of four reference samples (labeled A, B, C, and D). The RNA-seq methods were run using approximately 25 million reads per sample, obtaining estimates that were then compared to qPCR in Figures 1a and 1b, and the known ERCC spike-in mixtures in Figure 1c. In each of these we find that Isolator produces the highest correlation of the methods compared, though sometimes by a small margin. Despite two different technologies, and two different sets of genes, the qPCR benchmarks agree closely, ranking the methods nearly identically. Cufflinks, eXpress, Salmon, and RSEM/ML typically perform similarly to each other, with BitSeq, Kallisto, Sailfish, and RSEM/PM trailing slightly. Results are similar with ERCC spike-in controls, the notable exceptions being Kallisto and Sailfish, which show higher accuracy, and eXpress, which shows lower accuracy compared to their relative rankings in the qPCR benchmarks.

### 2.2 Estimate coherence

The four SEQC samples consist of two commercially available reference RNA samples, labeled *A* and *B*, and two synthetic samples formed by mixing *A* and *B* in specific proportions. Sample *C* is composed of 75% *A* and 25% *B*, while sample *D* is 25% *A* and 75% *B*. If we estimate transcript abundance for each sample, producing estimates *a, b, c, d*, then for these estimates to be in agreement (“coherent”, in our terminology) we should expect *c* ≈ 0.25*a*+0.75*b*, and similarly *d* ≈ 0.75*a* + 0.25*b*. In other words, mixing RNA then estimating transcript abundance should be approximated by estimating transcript abundance then mixing estimates.

To capture the notion of coherence, we measured the closeness of this approximation using proportionality correlation (Figure 1d).

Results from RSEM are particularly informative. We generated maximum likelihood and posterior mean estimates using the same aligned reads. Merely switching the estimate from the former to the latter increased the correlation by approximately 5%. This stands in contrast to comparisons to qPCR which suggest reduced accuracy of posterior mean estimates, demonstrating that coherence and accuracy, while related, are two separate axis of comparison.

Estimates made at the gene level (Supplementary Table 2) agree with these results, but are of uniformly higher correlation and show a much smaller gap between posterior mean and maximum likelihood estimate. Methods reporting posterior mean estimates (Isolator, BitSeq, RSEM/PM) again show the highest correlation and are in very close agreement.

### 2.3 Batch effects

We further examined the question of estimate variance by comparing the same samples sequenced on different flow cells and at different sites, again using data from the SEQC. We used 10 samples from the larger SEQC dataset, each consisting of a single lane from two separate flowcells sequenced at five different sites, all on Illumina HiSeq 2000 instruments, and compared pairwise agreement between estimates from these samples (Figure 2a). In the interest of a more fair comparison, Isolator was run on each sample in isolation without sharing any information.

These correlations are somewhat smaller than those in Table 1d, largely because of shallower sequencing: in Table 1d five flowcell lanes were combined, totaling ~25 millions reads per sample, here only one lane (~5 million reads) was used. Nevertheless, we again see posterior mean estimates (as in Isolator, BitSeq, and RSEM/PM) with significantly higher agreement between pairs.

When accounting for batch effects, bias correction methods can have a significant impact. To examine the efficacy of bias correction, we disabled this functionality on those methods that support it, repeated the experiment, and measured the change in pairwise correlation (Figure 2b). We see that bias correction is largely beneficial for each method. Isolator, Salmon, and Cufflinks show similar improvements with bias correction, though in a few cases Cufflinks slightly decreases agreement. Kallisto show a consistent, but very slight improvement. BitSeq’s bias correction sometimes has a very positive effect, but other times a negative or even disastrous effect. Estimates from the “MAY 2” samples in particular had far worse agreement with other samples with bias correction enabled. Here bias correction has dramatic and detrimental effects on some number of transcripts, driving the correlation to negative numbers.

While this benchmark is informative, it should be considered a lower bound on the batch effects and bias found in real experiments: these samples were sequenced at different labs, but using the same instrument model (HiSeq 2000) and from libraries prepared centrally. The fact that technical effects were so clearly present even under such controlled circumstances suggests that reproducibility should not be taken for granted in RNA-Seq experiments, particularly when it comes to batch effects.

**Figure 1:**
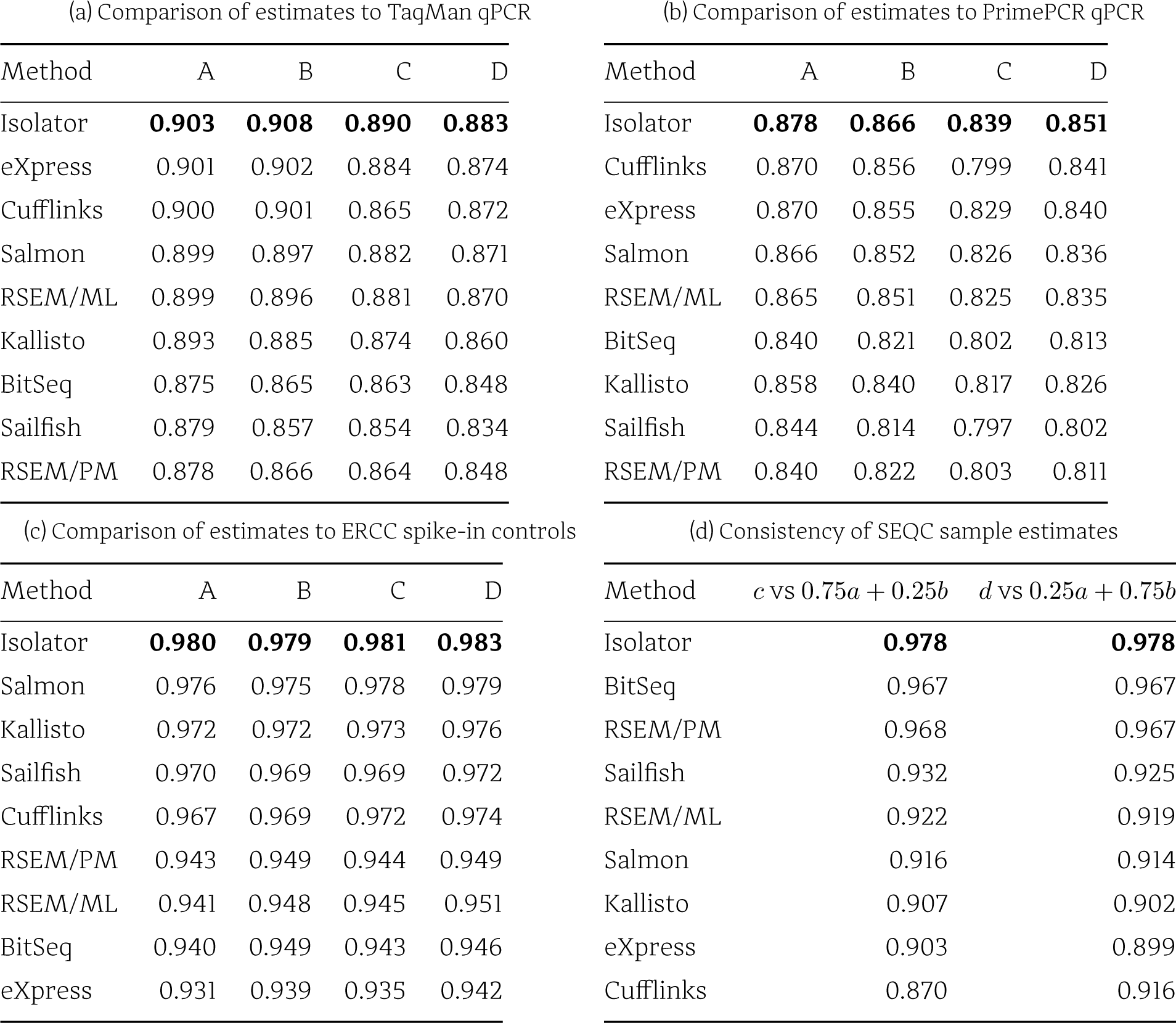
Benchmarks measuring the accuracy and coherence of estimates made from Sequencing Quality Control (SEQC) dataset using a variety of methods. All numbers are reported in proportionality correlation. (a) correlation between gene level quantification of 806 genes using TaqMan qPCR and RNA-Seq quantification. (b) correlation between gene level quantification of 18,353 genes using PrimePCR qPCR and RNA-Seq quantification. (c) correlation between known proportions of 92 ERCC spike-in controls and RNA-Seq quantification. (d) correlation between transcript-level estimates for the mixed SEQC samples C and D and weighted averages of estimates for A and B, corresponding to the mixture proportions for C and D. Sample C consists of 3/4 A and 1/4 B and sample D is 1/4 A and 3/4 B. Consistent estimates of transcript expression in A, B, C, and D should agree with these mixture proportions.

**Figure 2:**
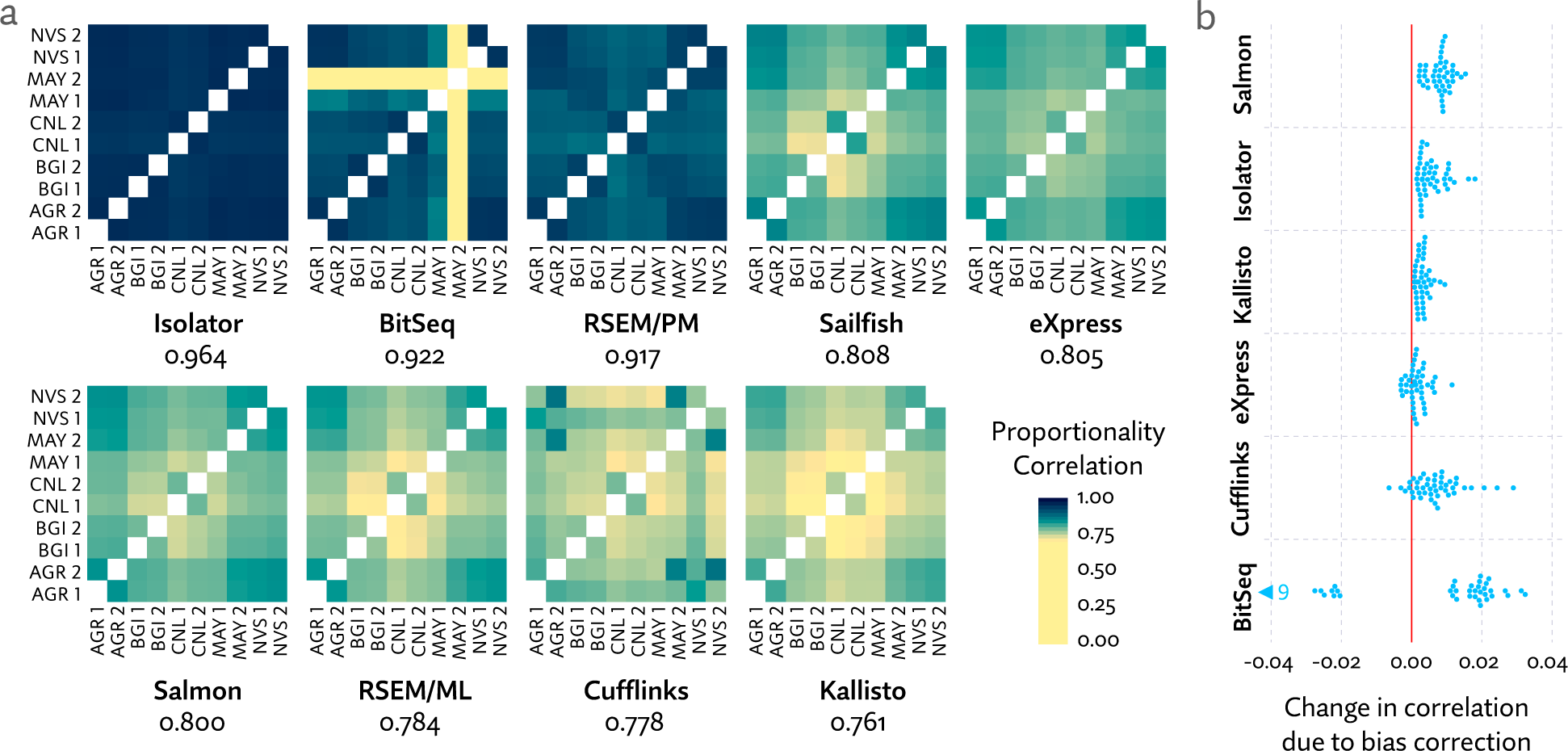
**a** A heatmap showing pairwise proportionality correlation between samples sequenced on two flowcells each at five sites, from centrally prepared libraries. Flowcells are numbered arbitrarily 1 or 2 and sequencing sites are abbreviated with three letter codes: Australian Genome Research Facility (AGR), Beijing Genome Institute (BGI), Cornell University (CNL), Mayo Clinic (MAY), and Novartis (NVS). Median proportionality correlation is listed below each heatmap. **b** The absolute change in correlation induced by enabling bias correction for methods that support it. For clarity this plot excludes points for BitSeq estimates of “MAY 2”, as bias correction has an extremely detrimental effect on these. Mean improvement in correlation was 0.008 with Salmon, 0.007 with Cufflinks, 0.006 with Isolator, 0.003 with Kallisto, 0.002 with eXpress, and -0.164 with BitSeq.

### 2.4 Accuracy in simulated data

To more directly demonstrate Isolator’s capacity to produce more accurate estimates by modeling entire experiments, we simulated a simple RNA-Seq experiment consisting of two conditions each with three replicates. Expression values were generated from a two-component log-normal model with parameters fit to Cufflinks estimates of data from Kuppusamy et al [10]. We generated 10 million, 100nt paired-end simulated RNA-Seq reads per sample using rlsim [23].

In this benchmark, Isolator significantly improves on existing methods (Table 1). It has the unique advantage of sharing information between replicates. With the assumption that genes tend to be similarly spliced between replicates, we are able to more effectively resolve transcript expression in complex loci. Kallisto, Salmon, RSEM/ML, and Cufflinks all perform very similarly. While comparisons to qPCR showed eXpress and Cufflinks with a significant advantage over RSEM/ML, Kallisto, and Bitseq, effective bias correction is less important in simulated data, plausibly explaining the difference. Although rlsim models some forms of bias, technical effects in RNA-Seq are not entirely understood, so these are likely understated in its model.

### 2.5 Agreement between two sequencing technologies

RNA-Seq reads become more informative with length and quantity. We compared data from the same sample sequenced twice, once with 300nt paired-end reads on a MiSeq sequencer and again with 100nt paired-end reads using a NextSeq 500, yielding approximately 4.5 million and 11 million reads, respectively. To elucidate the effect of deeper sequencing, we sampled without replacement smaller subsets of the 11 million NextSeq 500 reads and measured correlation between estimates made using these subsamples with estimates from the MiSeq data (Figure 3).

**Table 1:** Proportionality correlation between ground truth and estimates produced by each method using simulated RNA-Seq reads.

| Method | Correlation |
| --- | --- |
| Isolator | **0.919** |
| Kallisto | 0.887 |
| Salmon | 0.886 |
| RSEM/ML | 0.881 |
| Cufflinks | 0.881 |
| eXpress | 0.825 |
| Sailfish | 0.816 |
| RSEM/PM | 0.806 |
| BitSeq | 0.796 |

Isolator shows the highest correlation, but only when sequencing depth exceeds a few million reads (below which would be unusually shallow sequencing for an RNA-Seq experiment) where it overtakes eXpress by a small margin. The outliers in the this test are BitSeq and RSEM/PM. The former shows extremely high correlation with small numbers of reads, but contrary to all other methods, correlation rapidly decreases with sequencing deeper than 1 million reads. When posterior mean estimates are generated from a model using a uninformative prior, as is the case with BitSeq, transcripts with few or no reads may produce estimates that are highly influenced by transcript length, which plausibly explains this phenomenon. Yet, RSEM/PM does increase in correlation monotonically with sequencing depth, though the correlation remains very low. Even using all 11 million reads, RSEM/PM had a correlation of only 0.567 (data not shown).

**Figure 3:**
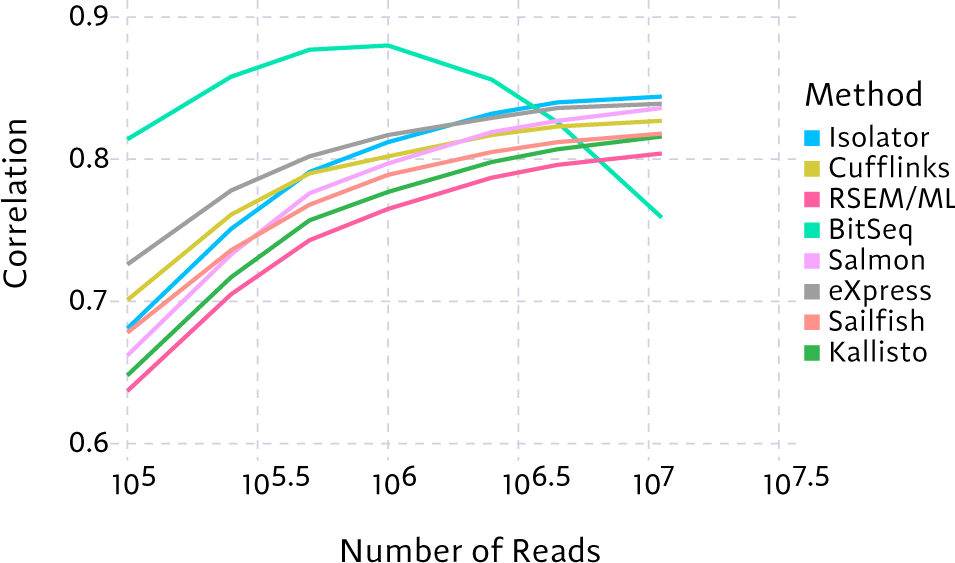
Proportionality correlation between estimates from 4.5 million 300nt MiSeq reads and progressively larger numbers of NextSeq 500 reads. For clarity, correlations for RSEM/PM are excluded as they were much lower than the other methods.

### 2.6 Case study: finding monotonic differential splicing

Splicing has been known to play an important role in developing hearts. Specific splicing changes during maturation have been shown to be conserved across chicken and mouse, and influenced by expression of CELF and MBNL [9]. Recently, stress-induced switching between two mutually exclusive isoforms of KHK has been shown to be driver of pathological cardiac hypertrophy [16].

We previously used Isolator to analyze splicing dynamics in human cardiomyocyte cells during maturation [10]. By modeling the entire RNA-Seq experiment and saving samples generated during sampling, Isolator is uniquely able to compute posterior probabilities corresponding to arbitrarily complex questions, within the confines of the model. As this experiment consisted of a number of samples from a number of conditions and time points, it did not easily lend itself to analysis using pairwise tests.

We instead searched for differential splicing events by computing the probability of “monotonic splicing”, which we defined as a splicing change that that is consistently observed to occur in the same direction between immature (H7 cells at day 20 and at day 30 infected with an empty vector) and mature samples (H7 1 Year, adult heart, and fetal atrium and ventricle). To compute the posterior probability of monotonic splicing, we simply count the number of samples from the model in which this pattern is observed, and divide by the total number of samples generated (350, in this case).

This approach identified 272 probable (posterior probability > 0.5) differential splicing events. We chose seven high probability candidates (ACOT9, IMMT, LTBP4, OBSL1, OSBPL1A, PALM, and TNNT2) to validate with qPCR (Supplementary Section 3.1), by measuring the expression of specific exons predicted to change in splicing. Five of these candidates (ACOT9, LTBP4, OBSL1, OSBPL1A, and TNNT2) showed a clear gradient between H7 day20, fetal, and adult tissue.

We analyzed the same data with the other seven methods. Exon-level fold change estimates generally agree between RNA-Seq methods, though far less so between RNA-Seq and qPCR, which could be caused by poor gene annotations or yet unidentified technical effects. Principal component analysis using transcript expression estimates also demonstrates some revealing differences (Supplementary Section 3.2).

### 2.7 Case study: novel mechanisms of cell fate specification identified by gene splice variants

Very little is known about the role of splice variants in mediating cell fate specification even though it has been shown that 92-95% of genes are differentially spliced in a cell or tissue specific manner [26, 20]. In order to identify novel mechanisms of cell fate specification in mesoderm development we began by analyzing RNA-seq data to determine differences in splice variants between human embryonic stem cell (hESC) derived cardiac progenitor cells (CPCs) and two endothelial lineages, hESC-derived endocardial endothelial cells (EECs) and hemogenic endothelial cells (HECs) as we have described previously [19]. 84 genes were identified as differentially spliced (defined here as a > 0.3 shift in relative transcript abundance with > 0.5 posterior probability) between these populations (Figure 4a). Principle component analysis indicated that splice variants, independent of gene expression levels, delineate endothelial populations relative to the CPCs (Figure 4b). Furthermore, these data indicate that splice variants alone are sufficient to differentiate these closely related lineages. Gene ontology analysis of the 84 differentially spliced genes showed appropriate enrichment for categories associated with alternative splicing (Figure 4c). Upon closer examination, we identified neuropilin-1 (NRP1) as an excellent candidate molecule that showed highly distinct expression profiles of splice variants between the cardiac and hemogenic endothelial lineages (Figure 4d-f).

**Figure 4:**
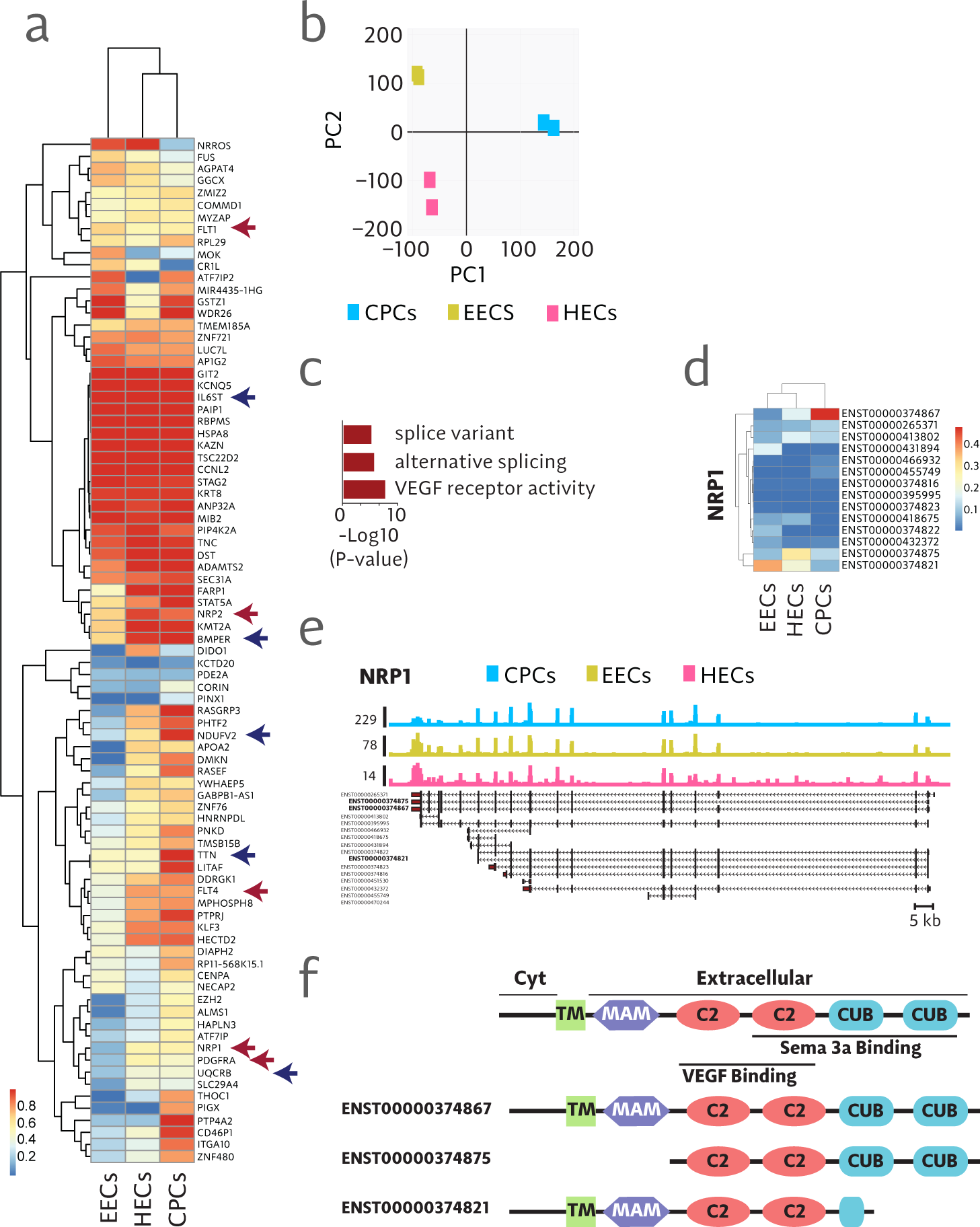
**a** Hierarchical clustering of the dominant splice variant for 84 differentially spliced genes. Blue arrows indicate genes with known function in endothelial or heart development. Red arrows indicate genes in the class of receptors that bind VEGF. **b** Two dimensional principle component analysis in mesodermal progenitor populations showing discrete separation of populations entirely based on differences in splice variants. **c** Analysis of gene ontology showing enriched categories found in 84 differentially spliced genes. **d** Heat map showing expression of annotated isoforms of NRP1. **e** Read coverage over isoforms of NRP1. **f** Schematic diagram of protein domains encoded by splice variant variants of NRP1.

NRP1 plays a versatile set of roles involving regulation of signaling pathways that mediate angiogenesis, axon guidance, cell survival, migration, and cell invasion in a wide range of developmental states and tissue types [27]. It is a single pass transmembrane molecule that has multiple extracellular functional domains that interact with VEGF and/or semaphorin molecules [2]. NRP1 has been implicated in endothelial [11] and cardiac [5] development but the specific splice variants involved in mediating these processes has not been elucidated. Results shown here indicate that the cardiac splice variant contains all functional domains involved in binding VEGF and semaphorin 3a (Figure 4d-f). This is consistent with studies showing the requirement for NRP1 in mediating the activities of VEGF and semaphorins in cardiac development [5]. In contrast, we show that splice variants expressed in endothelial populations include truncated versions that either lack the semaphorin binding CUB domains and, in the case of HECs, express a secreted version of NRP1 (Figure 4d-f). This suggests that VEGF and not semaphorin 3a is the primary mechanism of signaling through NRP1 in endothelial development [11]. These results provide the first evidence to our knowledge that NRP1 splice variants may be key determinants of fate determination in mesoderm patterning.

## 3 Discussion

To assess the suitability of Isolator we have compared it to large number of alternative methods using a wide variety of benchmarks, with a particular focus on estimation variance and coherence. Data generated by the SEQC is ideally suited to examine this question, and indeed our analysis agrees with their results. The SEQC compared Cufflinks and BitSeq and found BitSeq to produce more consistent, less variable estimates. We took a closer look at this phenomenon, controlling for alignment method by using the same set of alignments for every method (excluding the alignment-free methods Salmon, Sailfish, and Kallisto), and comparing a wider variety of methods. In this broader context it becomes clear that this separation is a consequence of the choice in estimators. Unmoderated maximum likelihood methods, though often accurate (i.e. low bias), frequently demonstrate less consistency (i.e. high variance) than methods that use MCMC sampling and report posterior means or medians. This is underappreciated but should be unsurprising. Because of the ambiguity of sequencing data, the likelihood functions for relative expression are often nearly plateaus. Maximum likelihood estimates are thus precarious compared to posterior mean estimates. Neither estimate is inherently more correct, and robust analysis should make use of the full posterior distribution, but we argue that posterior mean or median estimates are more useful in practice because they tend to have lower variance.

Excluding Isolator, methods that report posterior mean estimates show seemingly poor accuracy. This is largely driven by low-expression transcripts. A transcript with no reads has a maximum likelihood expression of zero, but its likelihood is positive over a range of non-zero values, so the posterior mean must necessarily also be positive. With a flat, or “non-informative,” prior, expression values for these low-expression transcripts are often quite high and highly determined by transcript length. Isolator avoids this problem by using a model with informative priors: namely encoding assumptions that most transcripts have low expression and that expression and splicing tend to be similar between samples. In this way, it is able to avoid what seems like a tradeoff and produces estimates that are both relatively low bias and low variance.

Fully Bayesian methods like the one presented here possess a degree of subjectivity that sometimes give researchers pause. Although we have found the results from Isolator to be insensitive to precise values given to hyperparameters, they are chosen in advance and with a degree of arbitrariness. What is often ignored is that the alternative, methods based on unregularized maximum likelihood estimates, in practice often necessitate a more insidious form of subjectivity: contriving cutoffs to filter out unreliable point estimates of low expression genes or transcripts. While the priors in Bayesian methods are purely explicit, ad hoc filtering of unregularized estimates can often be a form of off-the-books subjectivity–critical to the results, yet unmentioned or only alluded to in manuscripts. In this paper we have described a method that effectively shares information between samples in RNA-Seq experiments while rigorously quantifying uncertainty, leading to more accurate results across a wide range of benchmarks. We believe this work indicates a natural direction towards more reliable and powerful studies of transcriptomics, which future work can follow by scaling this methodology to larger numbers of samples, allowing for more flexible models, and accounting for more forms of uncertainty and technical effects.

## 4 Online Methods

### 4.1 Hierarchical Modeling of RNA-Seq experiments

Input to Isolator consists of gene annotations in GTF/GFF format, one or more sorted BAM [13] file, and optionally a reference sequence, which if provided is used in bias correction.

To model RNA-Seq experiments we use a hierarchical model consisting of three levels: samples or replicates, conditions, and the experiment, with any number of conditions, and any number of replicates within those conditions. We model condition-wise and experiment-wise expression for each transcript using a Gamma distribution, parameterized by mean and shape. Shape, which capture biological variability, are shared across conditions.

We further parameterize the model to capture splicing. Despite the compositional nature, for efficiency purposes we use independent Normal distributions the model splicing rates at the condition and experiment level, where a splicing rate is an isoforms proportion relative to total expression of the its gene.

Parameters are updated in sequence using a Gibbs sampling approach. The output generated by Isolator consists of each sample of every parameter serialized in an HDF5-based format, along with basic diagnostic information. Point estimates, summary statistics, and posterior probabilities can then be calculated from this output using a secondary command (isolator summarize).

### 4.2 Efficient Sampling

The isoform quantification or deconvolution problem is most easily thought of as an additive mixture model in which the component distributions (the isoforms) are known, but the mixture coefficients (isoform abundances) must be inferred. The standard approach to this problem is either expectation maximization (EM) if the maximum likelihood solution is desired, or Gibbs sampling if, as we are, estimating the posterior distribution. We have taken a different approach and instead rely on slice sampling [17].

The trade-offs between slice sampling and Gibbs sampling for the isoform quantification problem are subtle. Subsequent samples drawn from a slice sampler are generally less autocorrelated than those drawn from a Gibbs sampler. As a result, fewer samples need to be generated to adequately explore a distribution. However, each of these samples is more computationally expensive to compute, so one might compensate for autocorrelation in a Gibbs sampler by drawing more samples in less time. It is not obvious, then, that one approach is inherently more efficient than the other.

Because the likelihood function in this problem is simple, and we sample from the posterior probability directly, without introducing latent variables, we are afforded some optimization opportunities not available in a Gibbs sampler. Specifically, we implement the likelihood function using SIMD (single instruction, multiple data) instructions, which are available nearly all CPUs from the last decade. This, in combination with some numerical approximations, allow us to compute the likelihood function over an order of magnitude faster, than a more straightforward implementation. Run time for Isolator is generally significantly less than that of other sampling based implementations, though alignment-free, maximum-likelihood methods like Salmon and Kallisto remain the most efficient methods (Supplementary Figure 1)

### 4.3 Correcting for Bias

Isolator attempts to model and correct for multiple factors that can conflate the transcript expression estimates.

Perhaps the most prominent source of nonuniformity is sequence specific bias present at fragment ends. A probable contributor of this bias is cDNA synthesis by random priming, which is part of popular RNA-Seq protocols such as Illumina’s TruSeq kit. We have previously published a model that successfully accounts for such effects by training a sparse graphical model [7].

Beyond this sequence context bias, there is residual fragment GC-bias, which may be an artifact of PCR amplification. We observe that fragments with very high or very low GC-content are less likely to be sequenced than expected, even after accounting for sequence bias. We model this by binning fragments by GC content, computing expected and observed frequencies, and adjusting by their ratio.

Selection of polyadenylated transcripts is a common step mRNA-Seq, used to avoid sequencing introns, partially degraded transcripts, and ribosomal RNA. This can cause enrichment of reads at the 3’ end of transcripts if only partial transcripts are captured. To model the effect we fit a one parameter model in which a transcript is truncated at any position with probability *p*. The probability of observing a fragment ending at a position *k*, counting from the 3’ end, is then proportional to the probability that the transcript was *not* truncated before *k*, which is (1 –*p*)^*k*^. When fit, *p* is typically quite small, on the order of 10^−5^, in which case this correction has little effect on transcripts shorter that several kilobases.

Lastly, subtle implications arise from random fragmentation steps in many protocols. Existing statistical models assume fragments are sampled uniformly at random from a transcript. However, this does not exactly match the implications of random fragmentation in which, under an ideal model, *breakpoints* rather than fragments are introduced uniformly at random. For a fragment to be observed it must pass size selection and have fallen between two breakpoints, or one breakpoint and the end of the transcript. Since the ends of a transcript act as fixed breakpoints, the result is some enrichment of fragments at either end of the transcript. We compute this enrichment exactly for various transcript length, and use interpolation to approximate the effect.

